# OMAmer: tree-driven and alignment-free protein assignment to subfamilies outperforms closest sequence approaches

**DOI:** 10.1101/2020.04.30.068296

**Authors:** Victor Rossier, Alex Warwick Vesztrocy, Marc Robinson-Rechavi, Christophe Dessimoz

**Affiliations:** Department of Computational Biology, University of Lausanne, Switzerland; Center for Integrative Genomics University of Lausanne, Switzerland; SIB Swiss Institute of Bioinformatics, Lausanne, Switzerland; Department of Ecology and Evolution, University of Lausanne, Switzerland; Department of Genetics, Evolution, and Environment, University College London, UK; Department of Computer Science, University College London, UK

## Abstract

Assigning new sequences to known protein families and subfamilies is a prerequisite for many functional, comparative and evolutionary genomics analyses. Such assignment is commonly achieved by looking for the closest sequence in a reference database, using a method such as BLAST. However, ignoring the gene phylogeny can be misleading because a query sequence does not necessarily belong to the same subfamily as its closest sequence. For example, a hemoglobin which branched out prior to the hemoglobin alpha/beta duplication could be closest to a hemoglobin alpha or beta sequence, whereas it is neither. To overcome this problem, phylogeny-driven tools have emerged but rely on gene trees, whose inference is computationally expensive.

Here, we first show that in multiple animal and plant datasets, 18 to 62% of assignments by closest sequence are misassigned, typically to an over-specific subfamily. Then, we introduce OMAmer, a novel alignment-free protein subfamily assignment method, which limits over-specific subfamily assignments and is suited to phylogenomic databases with thousands of genomes. OMAmer is based on an innovative method using evolutionarily-informed *k*-mers for alignment-free mapping to ancestral protein subfamilies. Whilst able to reject non-homologous family-level assignments, we show that OMAmer provides better and quicker subfamily-level assignments than approaches relying on the closest sequence, whether inferred exactly by Smith-Waterman or by the fast heuristic DIAMOND.

OMAmer is available from the Python Package Index (as omamer), with the source code and a precomputed database available at https://github.com/DessimozLab/omamer.

## Introduction

Assigning new sequences to known protein families is a prerequisite for many comparative and evolutionary analyses (Glover *et al*., 2019). Functional knowledge can also be transferred from reference to new sequences assigned in the same family (Gabaldón and Koonin, 2013).

However, when gene duplication events have resulted in multiple copies per species, multiple “subfamilies” are generated, which can make placing a protein sequence into the correct subfamily challenging. Gene subfamilies are nested gene families defined after duplication events and organized hierarchically into gene trees. For example, the epsilon and gamma hemoglobin subfamilies are defined at the placental level, and nested in the adult hemoglobin beta subfamily at the mammal level (Opazo *et al*., 2008). Both belong to the globin family that originated in the LUCA (last universal common ancestor of cellular life).

Gene subfamily assignment is commonly achieved by looking for the most similar sequence (or “closest sequence”, *see Discussion)* in a reference database, using a method such as BLAST or DIAMOND (Altschul *et al*., 1990; Buchfink *et al*., 2015), before assigning the query to the subfamily of the closest sequence identified. For example, EggNOG mapper uses reference subfamilies from EggNOG to functionally annotate millions of unknown proteins of genomes and metagenomes (Huerta-Cepas *et al*., 2017, 2019). Briefly, each query is assigned to the most specific gene subfamily of its closest sequence, inferred using DIAMOND, with functional annotations then transferred accordingly.

However, ignoring the protein family tree can be misleading because a query sequence does not necessarily belong to the same subfamily as its closest sequence (Fig. 1). For instance, if the query branched out from a fast evolving subtree, its closest sequence might not belong to that subtree, but to a more general subfamily, or even not be classifiable in any known subfamily (Fig. 1. B). Or, in case of asymmetric evolutionary rates between sister subfamilies, the closest sequence might belong to a different subfamily altogether (Fig. 1. C). The prospect of observing these two scenarios is sustained by the long-standing observation that duplicated proteins experience accelerated and often asymmetric evolution (Conant and Wolfe, 2008; Sémon and Wolfe, 2007).

**Fig. 1.**
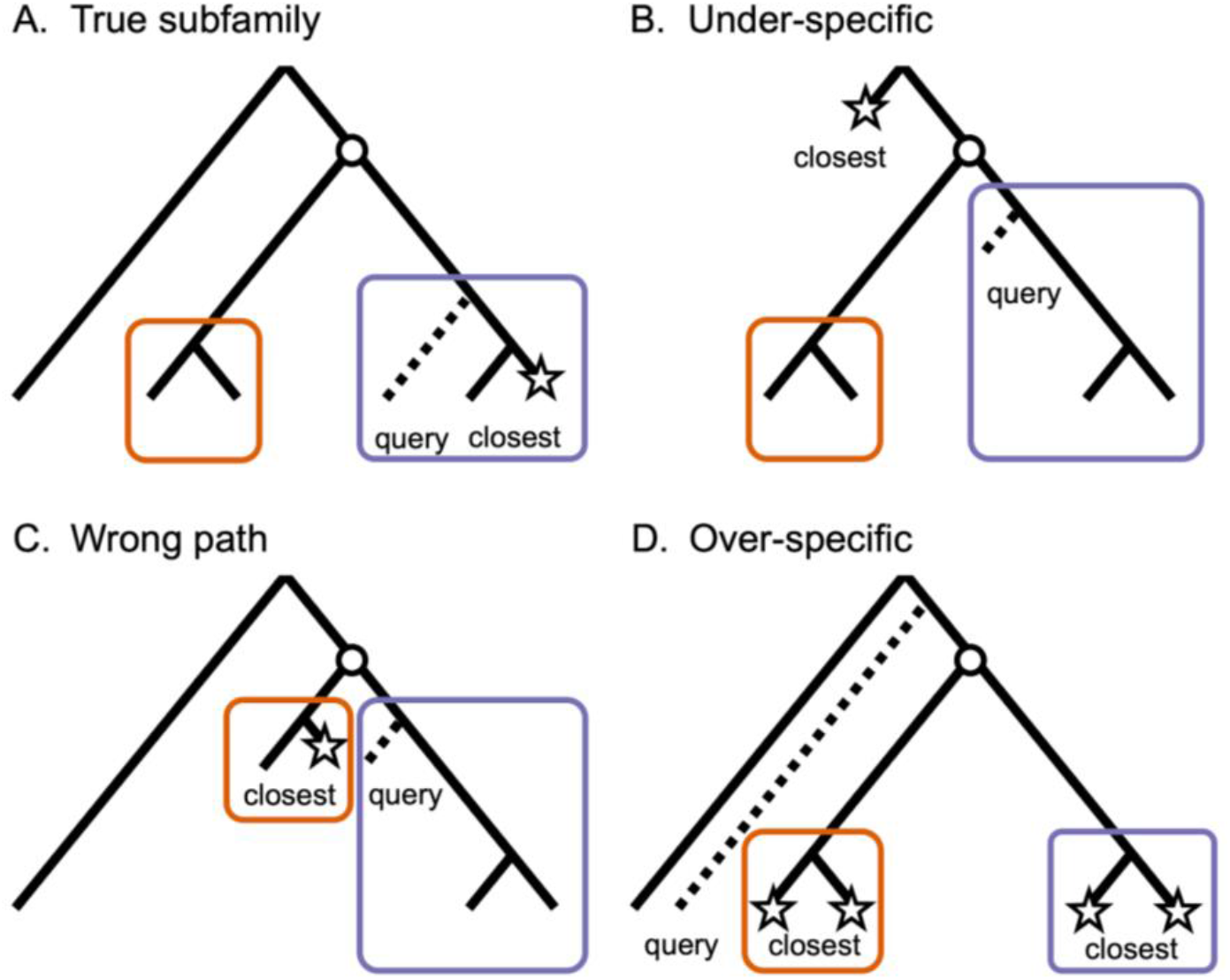
The closest sequence to a query does not necessarily belong to the same subfamily. This figure conceptualizes the four possible closest sequence locations relative to the query. On each tree, the true position of the query is indicated by a dashed branch, while its closest sequence(s) in the family is indicated by a star. The circle represents a duplication event leading to two subfamilies depicted as color boxes. Scenario A is the only one in which the closest sequence is in the same subfamily as the query. Note that for scenarios B and C to happen, the rate of evolution needs to vary across the tree (departure from a “molecular clock”), whereas scenario D can even happen under a uniform rate of evolution.

Moreover, the closest sequence to the query can belong to an over-specific subfamily even without any departure from the molecular clock in the family tree (Fig. 1. D). Such cases may occur when the query branched out before the emergence of more specific (nested) subfamilies. Indeed, all known proteins from the same clade as the query can belong to nested subfamilies. Moreover, even when not all proteins belong to such nested subfamilies, the closest sequence may still belong to an over-specific subfamily by chance. Since duplications are common in evolution (Conant and Wolfe, 2008), finding such nested subfamilies as close relatives to the query divergence is expected to be common. To avoid such errors, protein subfamily assignment tools relying on gene trees have been proposed (Schreiber *et al*., 2014; Tang *et al*., 2019). In short, these start by assigning queries to families with pairwise alignments against Hidden Markov profiles of reference families. Then, fine-grained assignments to subfamilies are performed with tree placement tools, which typically attempt to graft the query on every branch of the tree until maximizing a likelihood or parsimony score (Barbera *et al.,* 2018). However, gene tree inference is computationally expensive and therefore not scalable to the exponentially growing number of available sequences.

As a more scalable alternative to gene trees, the concept of hierarchical orthologous groups (HOGs) (Altenhoff *et al*., 2013) provides a precise definition of the intuitive notion of protein families and subfamilies. Each HOG is a group of proteins descending from a single speciation event and organized hierarchically. Moreover, they collectively provide the evolutionary history of protein families and subfamilies, like gene trees. While the oldest HOG in the family hierarchy (“root-HOG”) is the family itself, the other nested HOGs are its subfamilies. Thus, HOGs up to 100,000 members and covering thousands of species are available in large-scale phylogenomic databases (Altenhoff *et al*., 2018; Huerta-Cepas *et al*., 2019; Kriventseva *et al*., 2019).

Here, we first demonstrate on six animal and plant proteomes (sets of proteins from a given species, see *Methods)* that 18 to 62% of assignments by closest sequence go to incorrect, mostly over-specific, subfamilies. To overcome this problem, we introduce OMAmer, a novel alignment-free protein subfamily assignment method, which limits over-specific subfamily assignments and is suited to phylogenomic databases with thousands of genomes. We show that OMAmer is able to assign proteins to subfamilies more accurately than approaches relying on the closest sequence, whether inferred exactly by Smith-Waterman or by the fast heuristic DIAMOND. Furthermore, we show that by adopting efficient alignment-free *k*-mer based analyses pioneered by metagenomic taxonomic classifiers such as Kraken or RAPPAS (Wood and Salzberg, 2014; Linard *et al*., 2019), and adapting them to protein subfamily-level classification, OMAmer is computationally faster and more scalable than DIAMOND.

## Materials and methods

### The OMAmer algorithm

In this section, we describe the two main algorithmic steps which make OMAmer more precise and faster than closest sequence approaches. First, to speed-up the protein assignment step, OMAmer preprocesses reference hierarchical orthologous groups (HOGs) into a *k*-mer table (Fig. 2). For each *k*-mer and family (root-HOG), this table stores the subfamily (sub-HOG) where the *k*-mer has most likely arisen (the most specific HOG containing all occurrences of the given *k*-mer within the root-HOG). Then, these evolutionarily-informed *k*-mers are used to yield more precise subfamily assignments by reducing over-specific assignments (Fig. 3).

**Fig. 2.**
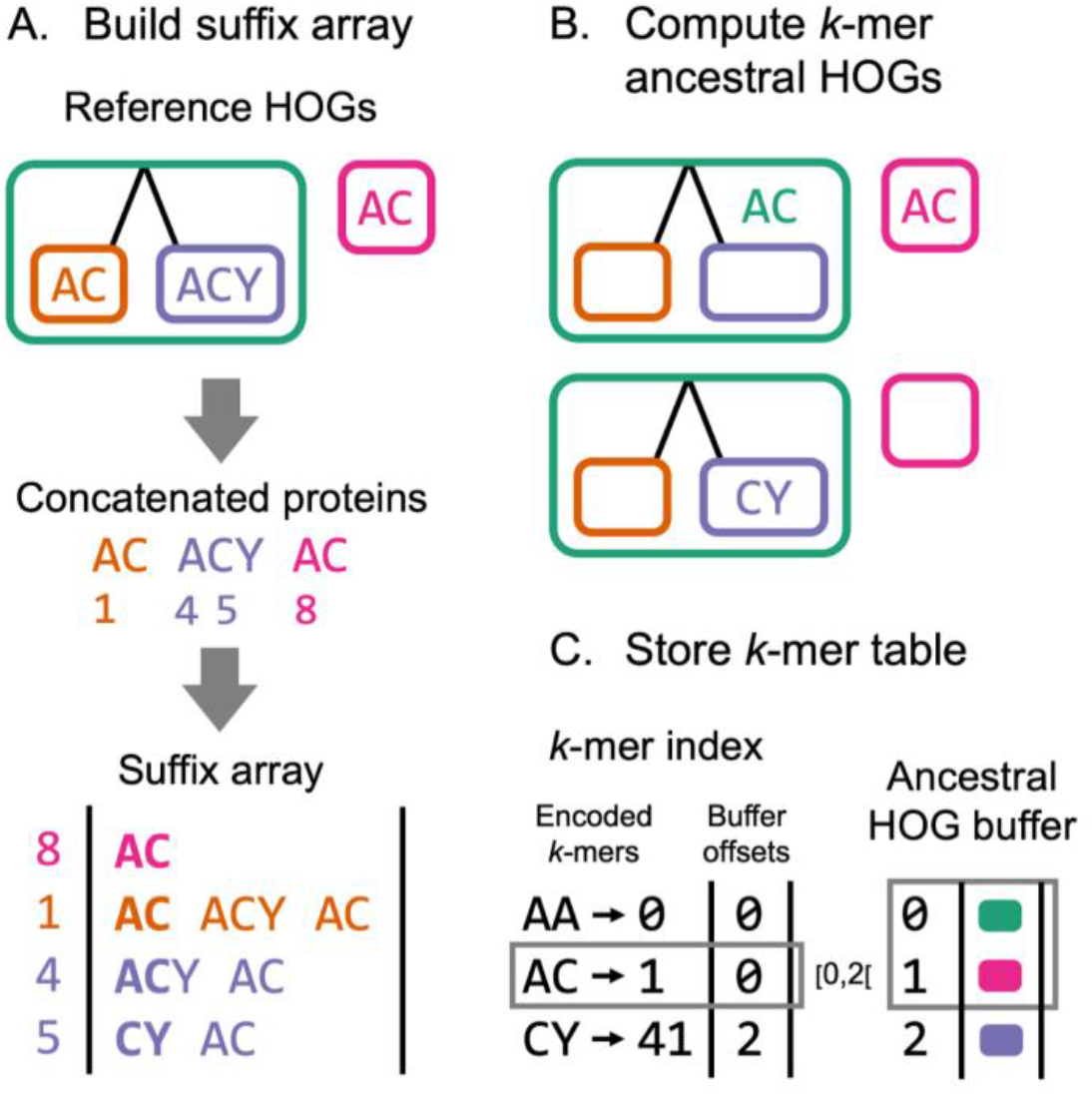
OMAmer algorithm for compact *k*-mer table precomputation. A. To efficiently preprocess the *k*-mer table, a suffix array is first built from concatenated protein sequences of reference hierarchical orthologous groups (HOGs), encoding families (root-HOGs) and subfamilies (sub-HOGs). Numbers indicate suffix offsets in the concatenated protein array and bold characters highlight *k*-mers at the beginning of suffixes. B. The *k*-mer ancestral HOG (where the *k*-mer has arisen) is approximated within each root-HOG as the last common ancestor among HOGs with the given k-mer. For example, since both the orange and purple sub-HOGs contain the “AC” *k*-mer, the ancestral HOG for that *k*-mer is the green root-HOG. C. The compact *k*-mer table includes an *k*-mer index mapping to a buffer that stores each *k*-mer ancestral HOGs. Note that each offset of the index corresponds to a k-mer integer encoding. As illustrated with the grey boxes, the “AC” *k*-mer (encoded as “1”) maps to the green and pink HOGs since these two lie within the [0,2[ offset interval in the buffer.

### *k*-mer table precomputation

To efficiently parse *k*-mer sets of reference HOGs, the suffix array (Manber and Myers, 1993) of all concatenated reference proteins is used as an intermediate data structure (Fig. 2. A.). There, all suffixes starting with a given *k*-mer are stored consecutively, which enables to quickly identify all HOGs containing the same k-mer using binary search.

Then, the *k*-mer ancestral HOG (where the *k*-mer has arisen) is approximated within each root-HOG as the last common ancestor (LCA) among HOGs containing the given *k*-mer (Fig. 2. B). Indeed, we assume that occurrences of the same *k*-mer in different members of a family mostly result from homology *(i.e.* same *k*-mer due to shared ancestry) rather than homoplasy *(i.e.* same *k*-mer arising independently). In the instances where the latter is true, the LCA approximation will favor overly general assignments. Thus, compared to the homoplasy assumption that would favor over-specific assignments, this approach is more conservative. Moreover, retaining a single ancestral HOG per *k*-mer and family reduces the memory footprint of the *k*-mer table.

Finally, to enable fast and memory efficient subfamily assignments, the resulting *k*-mer table is stored in the compressed sparse row format, consisting of two related arrays (Fig. 2. C). The *k*-mer index stores, at offsets corresponding to each *k*-mer integer encoding *(e.g.* 0 for AA, 1 for AC, etc.), offsets of the second array (the ancestral HOG buffer). There, the ancestral HOGs of each *k*-mer are stored consecutively. The formulae used to encode *k*-mers in integers is described in supplementary material.

### Family and subfamily protein assignment

The family (root-HOG) and subfamily (sub-HOG) protein assignment both rely on a common measure of similarity between the query protein and reference HOGs (the “OMAmer-score”). Essentially, this score captures the excess of similarity that is shared between the query and a given HOG, thus excluding the similarity with regions conserved in more ancestral HOGs. The OMAmer-score is computed in two main steps. First, a coarse alignment-free similarity is obtained by searching the *k*-mer table (Fig. 3. A). Second, this similarity is normalized to account for varying query lengths, composition biases, and sizes (number of different *k*-mers) of reference HOGs (Fig. 3. B). More details about the OMAmer-score computation are available in supplementary methods.

**Fig. 3.**
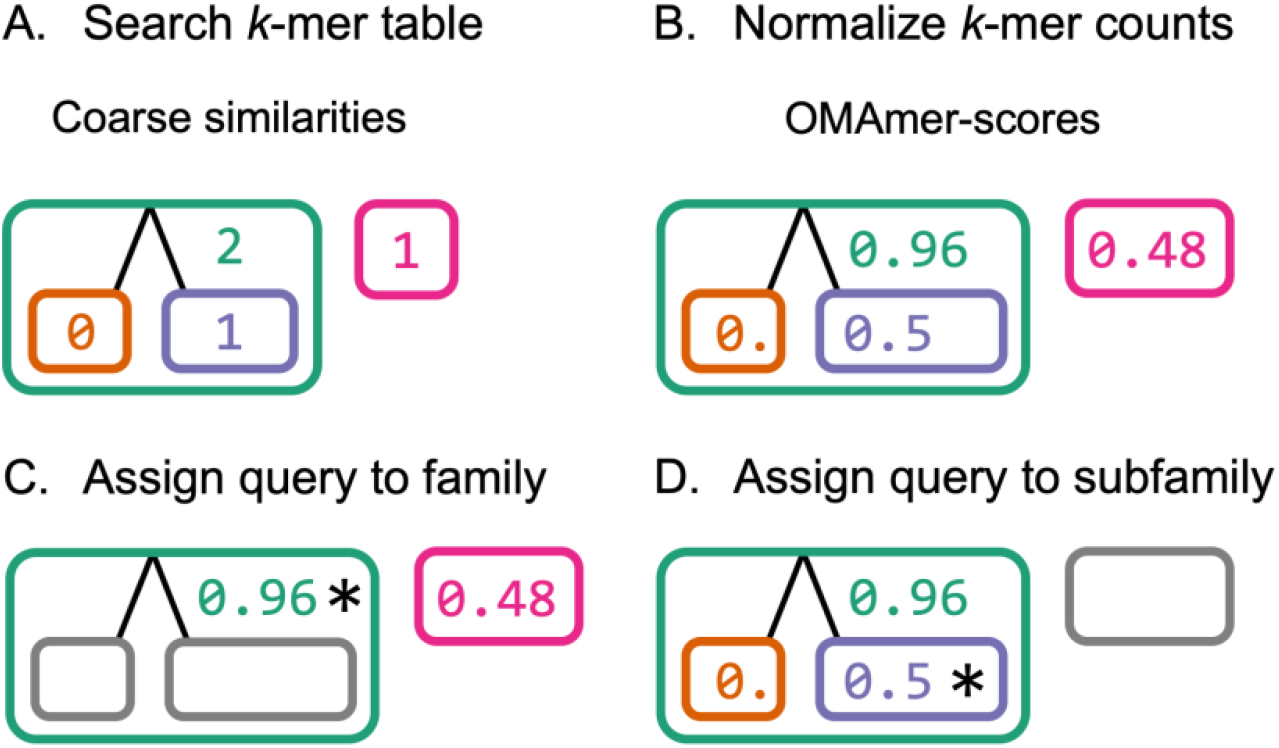
OMAmer algorithm for protein assignment to family and subfamily. A. Coarse alignment-free similarities are obtained by searching the k-mer table. B. These similarities are normalized into OMAmer-scores to account for varying query lengths, composition biases, and sizes of reference HOGs. C. The family (root-HOG) with the highest HOG-score is retained (shown with an asterisk) D. The assignment is refined to the most specific subfamily on the highest scoring root to leaf path (shown with an asterisk).

The protein is first assigned to the root-HOG with the highest OMAmer-score (Fig. 3. C). Indeed, at the family level, the OMAmer-score is analogous to other sequence similarity measures *(e.g.* alignment score) used to evaluate a probability of homology. Note that to speed-up the assignment, OMAmer-scores are computed only for the top 100 root-HOGs with the highest coarse alignment-free similarity. Then, the assignment is refined to the most specific sub-HOG on the highest scoring root-to-leaf path within the predicted root-HOG (Fig. 3. D). Indeed, at the subfamily level, OMAmer-scores are only comparable when descending from the same parent since they capture an excess of similarity relative to that parent.

Finally, to reduce the risk of false positive assignments, thresholds on the OMAmer-score can be applied at both steps. At the family level, this allows to avoid placing queries which have no homolog in the reference database. At the subfamily level, it penalizes more specific subfamilies to prevent over-specific assignments. Moreover, for applications where it is important to reject partial homologous matches (*e.g.* domain-level), OMAmer also outputs an “overlap-score” that measures the fraction of the query sequence overlapping with k-mers of reference root-HOGs (ignoring *k*-mers with multiple occurrences in the query sequence).

### Benchmarking

In this section, we describe the experiments conducted to evaluate the accuracy of OMAmer compared to closest sequence methods: Smith-Waterman (Smith and Waterman, 1981) and DIAMOND (Buchfink *et al*., 2015). Since placement in subfamilies initially requires accurate family-level assignments, we started by evaluating OMAmer at the family level *(i.e.* identifying the correct root-HOG). Second, to evaluate the impact of ignoring the phylogeny on subfamily assignments by closest sequences, we estimated the frequency of each closest sequence configuration (“true subfamily”, “under-specific”, “wrong-path” and “over-specific” [Fig. 1]). Third, we benchmarked subfamily-level assignments against closest sequence methods. Finally, we broke down the validation results of OMAmer by closest sequence configuration. The datasets and software parameters used in these experiments are described in supplementary methods.

### Family-level validation

Positive query sets were constructed as the sets of proteins from a given species contained in reference hierarchical orthologous groups (HOGs). We call these sets of proteins “proteomes” in this work. The proteins of that species were removed from the reference database used, before the *k*-mer index precomputation.

Since query proteins do not necessarily have homologous counterparts in the reference families (*e.g.* “orphan” genes, contamination, horizontal gene transfer), validating family assignments also required negative sets of non-homologous queries. Therefore, negative query sets were built with two approaches, while always matching the size of their corresponding positive set. In the first approach, random proteins were simply simulated with UniProtKB amino acid frequencies (release 2020_01) (UniProt Consortium, 2019) and sequence lengths of positive queries. The second approach was designed to resemble events of contamination or of horizontal gene transfer. Each negative query was randomly selected from a unique clade-specific family lying outside the taxonomic scope of reference families. In practice, clade-specific families were randomly selected among HOGs without parent (root-HOGs) at a given taxonomic level.

The resulting family assignments were compared with the truth set, and classified into true positives (TPs), false negatives (FNs) and false positives (FPs) for various score thresholds. FPs included negative queries assigned to a family as well as positive queries assigned to the wrong family (their relative proportion is shown in Supp. Fig. 2). The remaining positive queries were divided into TPs and FNs depending on whether the score for their family of origin passed the threshold, or not. Finally, precision, recall and accuracy (F1) were computed from TPs, FNs and FPs (Supp. Table. 1), defined according to the score threshold.

In the following experiments, to assess subfamily-level assignment separately from family-level assignment, we focused on the query sequences assigned to the correct family (*i.e.* the set of TPs at the threshold where F1 is maximal [F1_max_] for family assignment). Moreover, non-overlapping family-level TPs between methods being compared were further filtered out (sets of overlapping TPs are shown in Supp. Fig. 1.).

### Quantification of subfamily assignment errors by closest sequences

We used Smith-Waterman local alignments as reference to find the closest sequence (Smith and Waterman, 1981; Wolf and Koonin, 2012). Indeed, being an exact algorithm, Smith-Waterman is guaranteed to find the highest scoring match, and it is the standard approach in the field (Wolf and Koonin, 2012). Then, we classified each query according to the location of its closest sequence (Fig. 1.) as follows: a “true subfamily” configuration arises when the most specific HOG of the closest sequence is the same as the query one. An “over-specific” configuration arises when the most specific HOG of the query is ancestral to the most specific HOG of the closest sequence. Conversely, an “under-specific” configuration arises when the most specific HOG of the closest sequence is ancestral to that of the query. The last case is the “wrong-path” configuration, in which the most specific HOG of the query and of the closest sequence are in different parts of the family tree.

### Subfamily-level validation

To assess TPs, FNs and FPs at this level we took the view that an assignment to a subfamily also implies assignment to its “parental” subfamilies (if there are any). For instance, let us consider a nested gene family of alcohol dehydrogenases. Under this view, an assignment to the specific “alcohol dehydrogenase 1C” is also implicitly an assignment to “alcohol dehydrogenase 1”, as well as to “alcohol dehydrogenase”. In this case, if a method incorrectly assigns the protein to the subfamily “alcohol dehydrogenase 1B”, in addition to counting a FP (the gene is not a true member of subfamily “B”) and a FN (the gene is missing from subfamily “C”), we also count one TP for correctly assigning to the parental sub-HOG “alcohol dehydrogenase 1”. In effect, the prediction is regarded as being only partially wrong. Note that there is no TP counted for correctly implying an assignment to the root-HOG (alcohol dehydrogenase), because the present analysis only seeks to assess within-family placement.

In addition, we repeated the analyses using a second approach taking the more stringent view that there are no implicit predictions of parental subfamilies, therefore no reward is given for partial correctness. Thus, in the previous example, there would be no TP counted—only one FP and one FN. For both validation approaches, precision, recall and accuracy (F1) were computed from TPs, FPs, and FNs using the same formulae as at the family-level (Supp. Table 1).

### Performance experiments

To benchmark the computational performance of OMAmer and DIAMOND, we measured real and CPU time, as well as the maximum resident set size (memory) using the GNU time command. All timing was performed on machines containing identical hardware (dual-socket Intel Xeon E5-2680, 64GB of RAM). Single threaded versions of both methods were used, with timing repeated 10 times in order to ensure stability.

Databases of increasing size (20 to 200 proteomes, in steps of 20) were generated from Metazoan proteomes, with each including all of the previous and an extra 20 randomly selected species. The full proteomes of the initial 20 were used to query the databases of increasing size in order to gauge the scaling characteristics.

## Results

We first consider the problem of sequence placement at the overall family level (*i.e.* identifying the correct root hierarchical orthologous group, or “root-HOG”, defined at either *Metazoa* or *Viridiplantae).* Then, we present our analyses of the subfamily placement problem in four parts: First, we quantify the different types of errors resulting from the closest sequence criterion. Second, we show that OMAmer overcomes many of these errors, resulting in higher accuracy than closest sequence approaches. Third, we show that this accuracy improvement is mainly achieved by avoiding over-specific sequence classification. And fourth, we compare the computational cost and scaling of OMAmer and DIAMOND.

### At the overall family level, sequence placement is highly accurate

Query sequences must first be assigned to families before being placed within subfamilies. We evaluated this using DIAMOND and OMAmer, assessing the ability of the methods to either place a protein in its correct family, or to avoid placing a sequence with no homolog in the reference database (see *Methods*).

Both methods delivered similar and highly accurate results in placing platypus, spotted gar, and plant proteins (F1_max_ > 0.9; Supp. Fig. 2). The methods did not perform as well on the amphioxus proteome (OMAmer F1_max_ = 0.81-0.84; DIAMOND F1_max_ = 0.86-0.88; Supp. Fig. 2), but this is an outgroup to all other chordates in OMA, with a divergence of 600 MY (Peterson and Eernisse, 2016) to the closest species sampled (*i.e.* all vertebrates and urochordates) and with high levels of polymorphism which can result in alleles being misannotated as paralogs (Putnam *et al*., 2008; Huang *et al*., 2017; Kajitani *et al*., 2019). Still, this first analysis indicates that, with reference proteomes within the same phylum, family-level protein assignments are highly accurate.

### The closest sequence to a query is often not in the same subfamily

For a large proportion of query sequences (18-62%), the closest counterpart (inferred as the highest scoring Smith-Waterman match, see *Methods)* belongs to a different subfamily (Fig. 4. A). In such cases, the closest sequence most often belongs to a more specific subfamily (14-55% of all queries). These results highlight the need to account for the gene tree, especially in the presence of many nested subfamilies. Solving this problem is the primary aim of OMAmer.

**Fig. 4.**
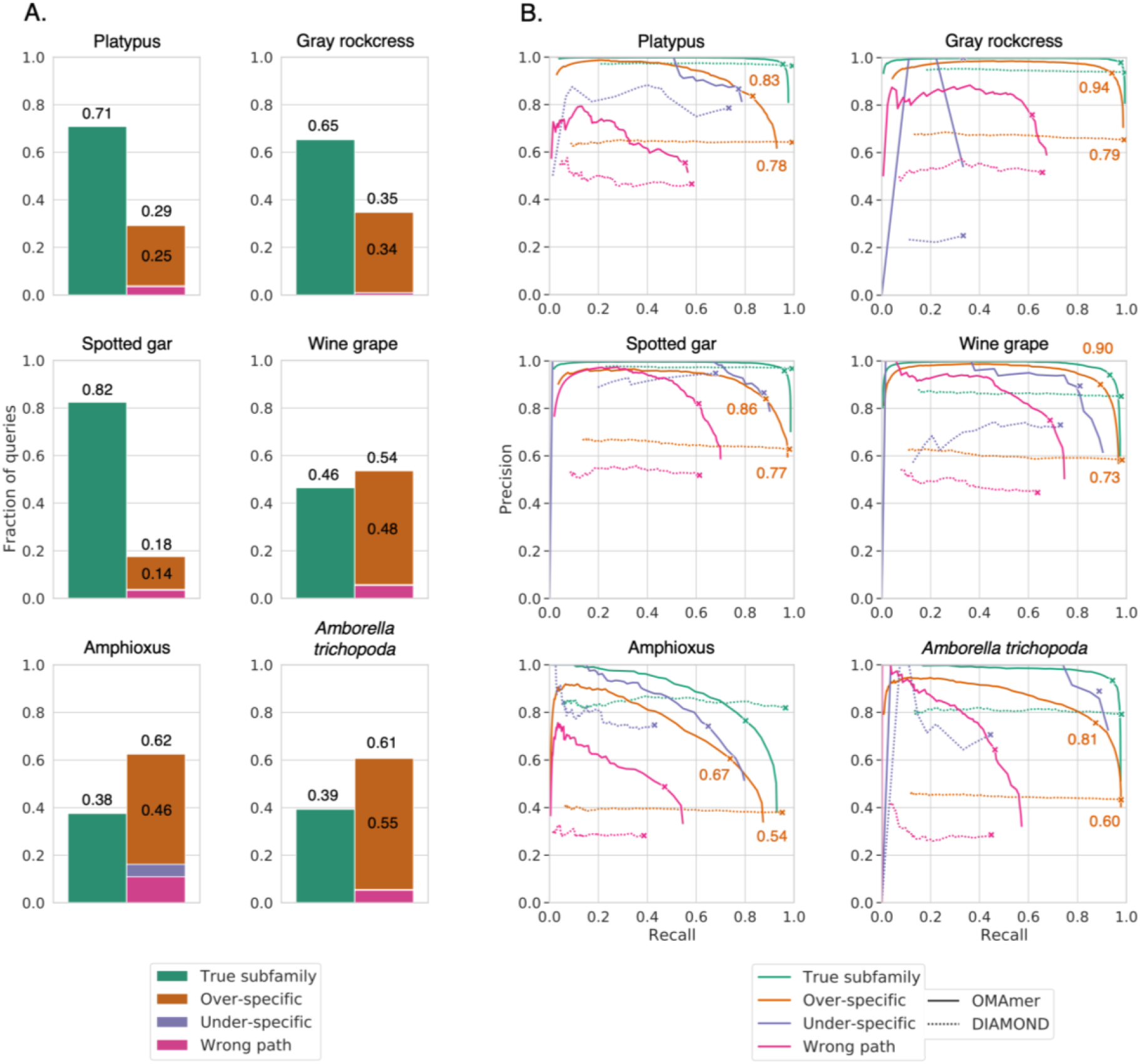
Frequency of closest sequence configurations defined in Fig. 1 and OMAmer accuracy for each. A. The closest sequence to a query was often found in another subfamily. Smith-Waterman alignments were used as proxies for closest sequences. B. “Over-specific” configurations were especially well dealt with by OMAmer. Each curve displays the range of trade-offs between precision and recall when varying the threshold on the OMAmer-score and on the DIAMOND E-value. They were computed by breaking down queries by closest sequence configurations as in panel A, before the validation procedure itself. Crosses indicate the location of F1_max_ values. “Over-specific” F1_max_ values are specifically annotated.

### OMAmer is more precise in subfamily placement

OMAmer systematically achieved, or equaled, the highest accuracy (F1_max_) across species (Fig. 5.). Specifically, increases in F1_max_ values between OMAmer and closest sequence methods ranged from 0.00 to 0.18. Moreover, OMAmer-score thresholds at F1_max_ were generally congruent (ranging from 0.10 to 0.16), although it was lower for amphioxus (0.06).

**Fig. 5.**
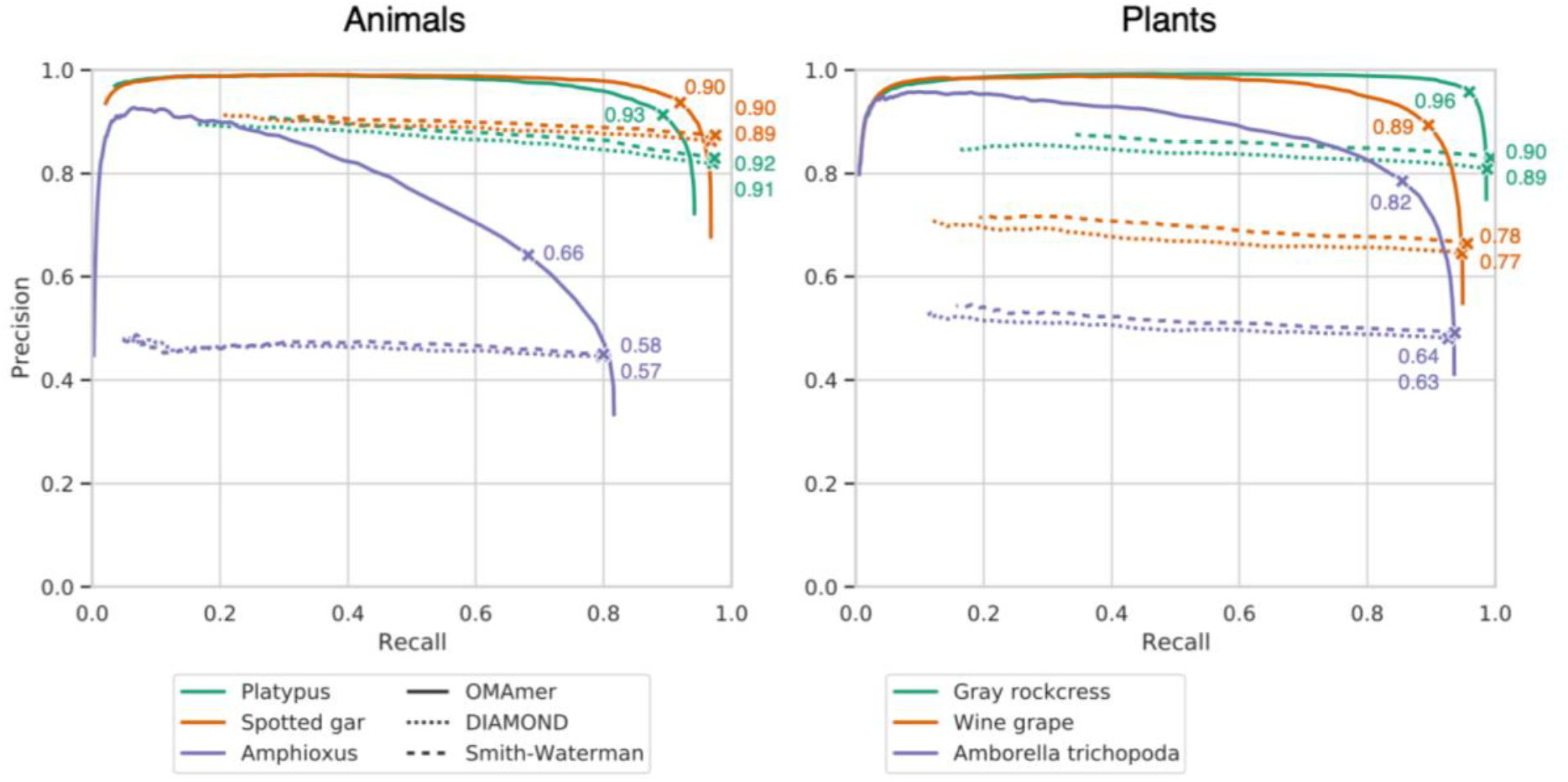
Comparison of subfamily assignments with OMAmer and by closest sequence (DIAMOND and Smith-Waterman). Each curve displays the range of trade-offs between precision and recall when varying the threshold either on the OMAmer-score, on the DIAMOND E-value or on the Smith-Waterman alignment score. F1_max_ values are indicated by crosses on each curve.

Importantly, OMAmer provides a genuine precision-recall trade-off, providing users with the possibility of obtaining very high precision, at the cost of lower recall. There is no such possibility with closest sequence methods: varying the E-value and alignment-score thresholds has very limited impact on precision (Fig. 5). These results are consistent with a second and more stringent validation procedure that does not reward assignments to correct parental subfamilies (Supp. Fig. 3).

### OMAmer deals especially well with over-specific closest sequences

As previously shown, over-specific placement is the most frequent mistake when only relying on assignments by closest sequences (Fig. 4. A). Since OMAmer was specifically designed to deal with such cases using evolutionarily-informed *k*-mers mapping toward ancestral subfamilies, we investigated whether this feature would explain OMAmer performance. Therefore, we reproduced the subfamily-level validation procedure with queries partitioned between the types of closest sequence configuration (“true subfamily”, “under-specific”, “wrong path” and “over-specific”) depicted in Fig. 1. and quantified in Fig. 4 A.

As expected, OMAmer was systematically more accurate than DIAMOND for queries in the “overspecific” configuration (Fig. 4. B). Specifically, for these queries, increases in F1_max_ values between OMAmer and DIAMOND ranged from 0.05 to 0.21. Moreover, OMAmer displayed a proportion of over-specific assignments (defined at F1_max_) 0.07 to 0.37 lower than Smith-Waterman and DIAMOND (Supp. Fig. 5). In animals, this performance for queries in the “over-specific” configuration was achieved while sacrificing very little accuracy for queries in the “true subfamily” configuration (from 0.02 to 0.11, Fig. 4. B). In plants, OMAmer remained more accurate even for queries in the “true subfamily” configuration, with increases in F1_max_ values between OMAmer and DIAMOND that ranged from 0.02 to 0.06. Queries in the “wrong-path” configuration were also placed more accurately by OMAmer, despite their small number. Finally, there were too few “under-specific” configurations to draw any conclusion.

Since DIAMOND is a closest sequence approach, like Smith-Waterman, it was expected to obtain precision values close to zero for queries in the “under-specific”, “wrong path” and “over-specific” scenarios. However, this behaviour is not observed here because the validation procedure rewards assignments in correct parental subfamilies even when the predicted exact subfamily is incorrect. By contrast, the more stringent validation procedure that does not reward assignments to correct parental subfamilies does yield precision values close to zero (Supp. Fig. 4. B). Apart from this difference, the results of this section are consistent between the two validation procedures (Supp. Fig. 4. B and 5).

The occasional and counterintuitive positive correlation between precision and recall that can be observed with OMAmer at low recall values, seemed to appear only when a few FPs subfamilies remained predicted at high OMAmer-score thresholds, while the number of TPs was steadily decreasing. Taking the example of the “wrong-path” Spotted gar proteins, 4 out of the 14 predicted subfamilies are FPs at recall of 0.02 obtained with the highest threshold value (0.99).

### OMAmer run time scales better than DIAMOND with the number of reference proteomes

In an empirical scaling analysis, we varied the number of reference proteomes in the database whilst querying a number of full-proteomes (see *Methods* for details). OMAmer achieved better scaling than DIAMOND in terms of CPU and real time when increasing the number of reference proteomes in the database (Fig. 6, left). Both methods, however, exhibited a similar increase in maximum memory usage (Fig. 6, center), with OMAmer initially using over 2GB and DIAMOND using less than 256MB on a database of 20 reference proteomes. In order to achieve this performance, OMAmer only stores *k*-mers once per root-HOG. This does require extra computation, with the overhead being reflected in its memory usage and time to build the database (Supp. Fig. 6), taking between 15-20 minutes in comparison to 1-2 minutes for DIAMOND.

**Fig. 6.**
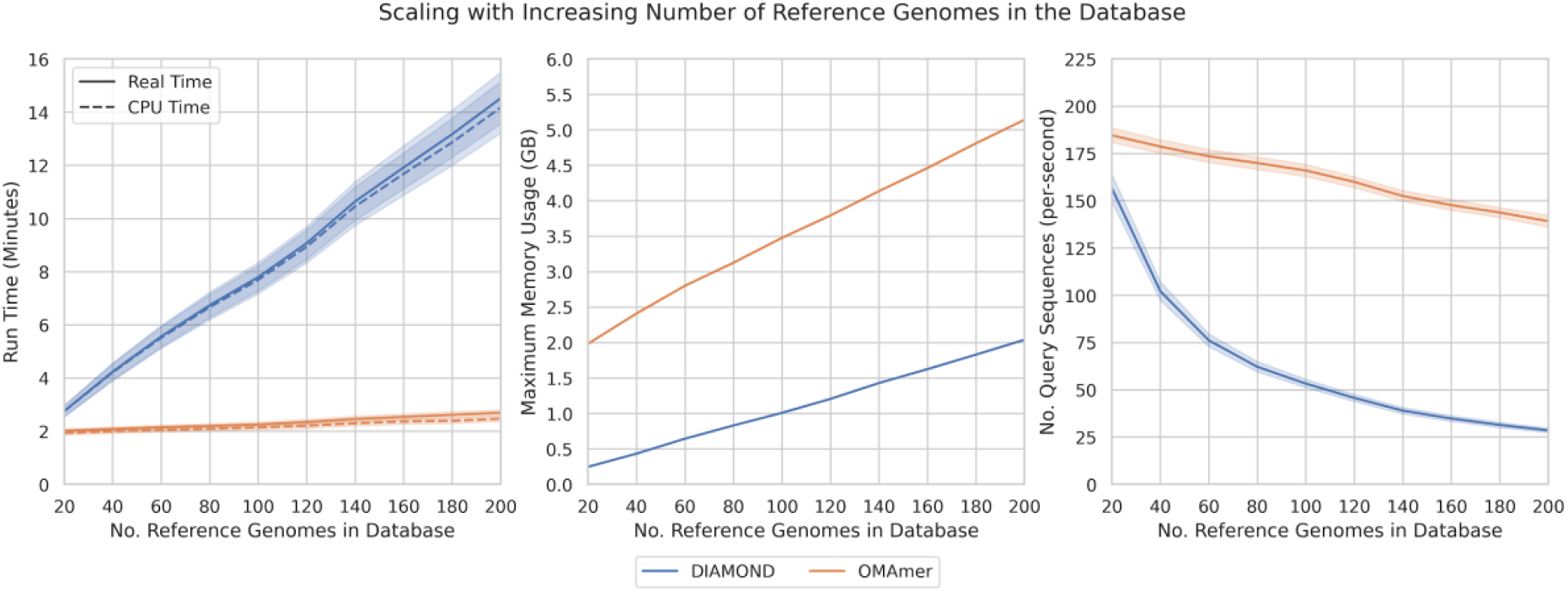
Comparison of the computational performance of family and subfamily assignments between OMAmer and DIAMOND. OMAmer scales better than DIAMOND in terms of real and CPU time (total time on the left, and sequence/second on the right), but requires somewhat more memory (center) Error bars shown for 95% confidence interval, estimated using 10,000 bootstraps.

To put the timing into context, OMAmer is processing about 150 query sequences per second (Fig. 6, right). DIAMOND starts with a similar performance, before trailing off to less than 30 with the largest number of reference proteomes.

## Discussion

In this study, we demonstrate that considering the phylogenetic relations between orthologous groups is essential for the problem of subfamily assignment. Indeed, although alignment-free, OMAmer generally outperforms closest sequence approaches, even when inferred by the exact SmithWaterman algorithm. In particular, OMAmer systematically equaled or out-performed Smith-Waterman for the best precision-recall trade-off (F1_max_).

However, the main advantage of OMAmer is its control over assignment precision through the setting of specific OMAmer-score thresholds that refrain over-specific placements. By contrast, relying on the closest sequence does not provide the ability for any precision-recall trade-off. Each assignment is bound to the most specific subfamily of the closest sequence, and varying the E-value threshold has a large impact on recall but almost none on precision. Thus, while closest sequence approaches are useful for cases where high recall is the overriding priority, OMAmer is more flexible and applicable in a broad range of contexts.

In addition to providing robust subfamily assignments, OMAmer scales better than DIAMOND, in terms of run time, with the number of reference proteomes. This is achieved with alignment-free sequence comparisons against hierarchical orthologous groups (HOGs) instead of approximate alignments against protein sequences. Indeed, in addition to removing the computational burden of sequence alignment, merging sequence information in HOGs drastically reduces the number of comparisons. This is especially true since the number of reference HOGs increases more slowly than proteins with the number of reference proteomes.

Large-scale sequencing projects of genomes or metagenomes add difficulties such as chimeric assemblies or contaminations, thus mixing gene families from different species. OMAmer was designed as a starting point for the integration of such heterogeneous data. Thus, instead of constraining subfamily assignments along the known taxonomy of query proteomes, OMAmer performs taxonomically blind assignments. We hope that this feature will enable diverse applications of OMAmer. For example, the detection of contamination and horizontal gene transfers could be achieved by including all kingdoms in the OMAmer database and searching for incongruent placement regarding the query taxonomy. In particular, confidence measures similar to the “Alien index” (Gladyshev *et al*., 2008) could be computed by subtracting the OMAmer-score of the highest-scoring taxonomically congruent HOG from the overall highest OMAmer-score potentially derived from a contaminant sequence. Other promising applications are the binning of protein-level metagenomic assemblies (Steinegger *et al*., 2019), and with some algorithmic adaptations, directly placing reads to skip genome assembly and annotation.

The OMAmer algorithm builds upon some key ideas of the metagenomic software Kraken, which classifies reads into the species taxonomy (Wood and Salzberg, 2014). Indeed, this task is analogous to protein subfamily assignments for two reasons. First, some prior knowledge, shaped as labelled reference sequences, is preprocessed before the assignment itself. Second, this prior knowledge is organized hierarchically in a tree graph. Thus, instead of relying on closest sequences, such methods of taxonomic classification exploit semi-phylogenetic information to improve their predictions. While MEGAN introduced the key idea of taking the LCA taxon among significant BLAST hits (Huson *et al.,* 2007), Kraken scaled up the approach by preprocessing LCA taxa in a database of taxonomically-informed *k*-mers (Wood and Salzberg, 2014).

While inspired by Kraken, the OMAmer algorithm features three key algorithmic innovations to fit the case of assigning proteins to subfamilies. The first difference lies in the types of events used to define clades or subtrees. Indeed, while taxa are defined by speciation nodes in Kraken, subfamilies are defined by duplication nodes in OMAmer. This is an important difference because duplication patterns are variable across protein families, whereas the reference taxonomy is the same for different genes and genomes in Kraken. Second, the dual problem of first placing sequences within families, followed by subfamily-level assignment is specific to OMAmer. Third, while Kraken relies on an arbitrary cut-off of one *k*-mer to avoid over-specific placements, OMAmer applies a user-defined threshold on the more refined OMAmer-score.

Beside closest sequence approaches, alignments to Hidden Markov Models (HMMs) have been extensively used for sequence to family or subfamily comparisons with tools such as HMMER3 (El-Gebali *et al*., 2019; Mi *et al*., 2019; Huerta-Cepas *et al*., 2019; Ebersberger *et al*., 2009). However, the use of HMMs is revealing a lack of scalability to phylogenomic database size. For instance, the developers of the EggNOG database reported that DIAMOND is considerably faster and achieves similar results to HMMER3, and have discontinued the use of HMMs in the latest EggNOG mapper release (Huerta-Cepas *et al*., 2017, 2019). Moreover, maintaining subfamily HMM models can be problematic because it relies on ad-hoc criteria for subfamily delineation *(e.g.* curated, family-specific E-value thresholds in Pfam (El-Gebali *et al*., 2019)). Finally, HMMs are tailored to detect remote homology rather than discriminating between specific subfamilies. Although this has benefited from hierarchically organized HMMs (Nguyen *et al*., 2016), the family breakdown is used to improve family assignments rather than finding specific subfamilies.

Due to the rapid emergence of alignment-free methods, covering various biological problems ranging from phylogenetic inference to metagenomic taxonomic profiling (reviewed in: (Zielezinski *et al.,* 2017)), the AFproject was launched to unite the benchmarking of these tools (Zielezinski *et al*., 2019). However, the available datasets to benchmark protein sequence classification in that project are organized according to the SCOPE database (Fox *et al*., 2014). There, each hierarchical level is either based on a degree of belief in homology among sets of proteins (families and superfamilies) or on structural similarities (folds and classes). By contrast, in this work, we seek to distinguish all subfamilies resulting from gene duplications, even recent ones yielding quite similar subfamilies. Of note, recent subfamilies can diverge in function (Naseeb *et al*., 2017) and thus be important for annotation.

In this work, we used the most similar sequence (whether inferred exactly by Smith-Waterman or by the fast heuristic DIAMOND) as reference to find the closest sequence. Although the highest scoring local alignment is not always the closest sequence in a phylogenetic sense (Koski and Golding, 2001), this is a commonly used approximation for classifying large numbers of orthologs (Li *et al*., 2003; Sonnhammer and Östlund, 2015; Huerta-Cepas *et al*., 2019) and has shown to give similar results in simulation (Dalquen *et al*., 2013) and empirical benchmarks (Altenhoff *et al*., 2016)

Although placing proteins at the overall family level appears to be easier than at the subfamily level, we start to see some degradation with the amphioxus sequences (last common ancestor to vertebrates 600MY [Peterson and Eernisse, 2016]). We expect further degradation for cases where query proteomes are even farther from the reference proteomes, because relying on *k*-mer exact matches is likely to be less sensitive than alignments such as provided by DIAMOND to detect distant homologs. Some avenues to increase OMAmer sensitivity in the absence of closely related reference species could be explored: the use of a reduced alphabet, which compresses the mutual information of sequences being compared (Edgar, 2004); or spaced seeds, *i.e.* non-contiguous *k*-mers, that have shown an increased sensitivity in metagenomics classification (Břinda *et al*., 2015). On the other hand, adding such very distant proteomes is expected to be much rarer than adding proteomes to an already sampled clade. This is especially true for the increase of sequences through projects such as i5k (insect genomes) (i5K Consortium, 2013) or the Vertebrate Genomes Project (Koepfli *et al*., 2015), where duplications and thus subfamilies are common and a solid backbone of reference proteomes are available. OMAmer is especially well positioned to help classify the genes from such projects, which will present a challenge for slower or less precise methods.

## Acknowledgements

We thank Adrian Altenhoff for fruitful discussions in the conception and validation of OMAmer and Yannis Nevers for proof-reading the manuscript. Computations were performed at the Vital-IT Center for high performance computing of the SIB Swiss Institute of Bioinformatics, as well as the Wally and Axiom clusters of the University of Lausanne.

## Supplementary material for OMAmer

### Supplementary methods

#### *k*-mer integer encoding

The integer encoding of a *k*-mer x formed of *k* numerical characters *x_i_*, ordered from *i* = 1 to *i = k*, from an alphabet *A* (A:0, C:1, …, Y:20) is defined as:

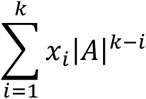

#### OMAmer-score computation

The coarse alignment-free similarity (1) is measured as the number of intersecting *k*-mers between the query *k*-mers *Q* and the HOG specific *k*-mers *H. H* includes *k*-mers specific to the HOG descendants but excludes the ones conserved in its ancestors. To compute (1), the number of intersecting *k*-mers between the query *k*-mers and each HOG ancestral *k*-mer set (the *k*-mers inferred to have arisen in the HOG) is retrieved from the precomputed *k*-mer table. Then, these counts are cumulated from leaves to root by adding the highest child HOG *k*-mer count to the current HOG count at each multifurcation.

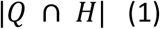

To account for the different sizes (number of different *k*-mers) of reference HOGs and the query composition bias, the expected number of shared *k*-mers between the query and the HOG observed in absence of homology, *i.e.* by chance, (2) is subtracted from (1) (3). OMAmer proposes a parametric (default OMAmer-score) and a non-parametric approach (sensitive OMAmer-score) to compute (2).

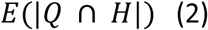

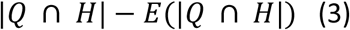

In the parametric approach, (2) is calculated as the number of query *k*-mers |*Q*| multiplied by the probability to observe one query *k*-mer *x_q_* in *H*.

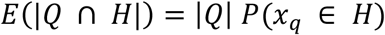

This probability is the inverse probability of not observing one *x_q_* in *H*.

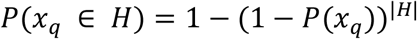

The probability of observing *x_q_* in a HOG of size one, *i.e.* with one *k*-mer, *P*(*x_q_*) is approximated as the mean frequency of query *k*-mers inside the *k*-mer table (the average fraction of HOGs containing each query *k*-mer *x_i_* [remember that each *k*-mer can only be stored once per root-HOG]).

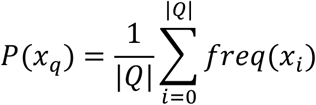

In the non-parametric approach, (2) is simply (1) obtained from a random permutation of the query sequence. In an attempt to conserve some local composition bias, the permutation is performed by shuffling windows of size six in addition to shuffling individual amino acids within each such window. Note that this approach additionally corrects for HOG composition biases.

Finally, to make the OMAmer-score comparable across queries, (3) is divided by |*Q*|, from which was subtracted the number of query *k*-mers shared with more ancestral HOGs.

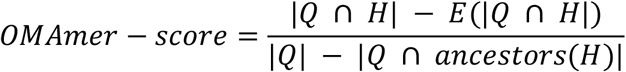

#### Datasets and software parameters

OMAmer was compared with two closest sequence methods lying at different extremes of the speedaccuracy tradeoff: DIAMOND (v0.9.14) and Smith-Waterman, respectively. Due to the computational cost of performing Smith-Waterman alignments, we used pre-computed alignments from OMA (January 2020) (Altenhoff *et al*., 2018). DIAMOND databases were built with default parameters, and searches for the most similar sequence were performed with effectively no significance requirement (E-value set to 1e6). The OMAmer *k*-mer table was built with a *k*-mer size of 6.

OMAmer directly yields family and subfamily predictions. For Smith-Waterman and DIAMOND, each query was assigned to the family and most specific subfamily of its closest reference protein. To obtain multiple precision-recall values, predictions were computed for multiple score thresholds: E-values of 1e-322 to 1e6 for DIAMOND, alignment scores of 1 to 5,000 for Smith-Waterman and OMAmer-scores of 0 to 0.99.

To make family-level assignments comparable and well differentiated from subfamily-assignments, we selected HOGs from OMA (January 2020) defined at the *Metazoa* and *Viridiplantae* taxonomic levels as root-HOGs (families), and their sub-HOGs as subfamilies. To avoid low-confidence families, we further filtered out root-HOGs with less than six proteins. We picked *Metazoa* because it is one of the largest clades in OMA and *Viridiplantae* due to the high number of duplications and thus subfamilies in this clade. Note, due to the addition of *Branchiostoma lanceolatum* in the January 2020 OMA release, we removed it from the reference database used (before the *k*-mer index precomputation) to keep the same evolutionary distance existing between *Branchiostoma floridae* and reference proteomes of the previous OMA release (June 2019).

Then, we selected six species as experiment targets picked because they stand as outgroups of large clades in OMA and thus display some variability in divergence ages to reference species. Platypus, spotted gar and amphioxus were selected in *Metazoa,* while Gray rockcress, wine grape and *Amborella trichopoda* were chosen in *Viridiplantae* (Supp. Table 2). Clade-specific root-HOGs used to build the negative query set were picked at the *Bacteria* taxonomic level.

The *Metazoa* reference dataset included 1,309,488 proteins from 201 species organized in 235,983 HOGs and including 12,178 root-HOGs. The *Viridiplantae* reference dataset included 554,389 proteins from 63 species organized in 304,838 HOGs and including 8,652 root-HOGs. The query datasets (proteomes) included 5,811, 7,7227,387, 6,239, 7,219 and 5,931 proteins of platypus, spotted gar, amphioxus, gray rockcress, wine grape and *Amborella trichopoda* species, respectively. 4,952, 6,308, 5,803, 5,261, 5,712 and 4,057 queries belonged to a sub-HOG in addition to the root-HOG.

## Supplementary figures

**Supp. Fig. 1.**
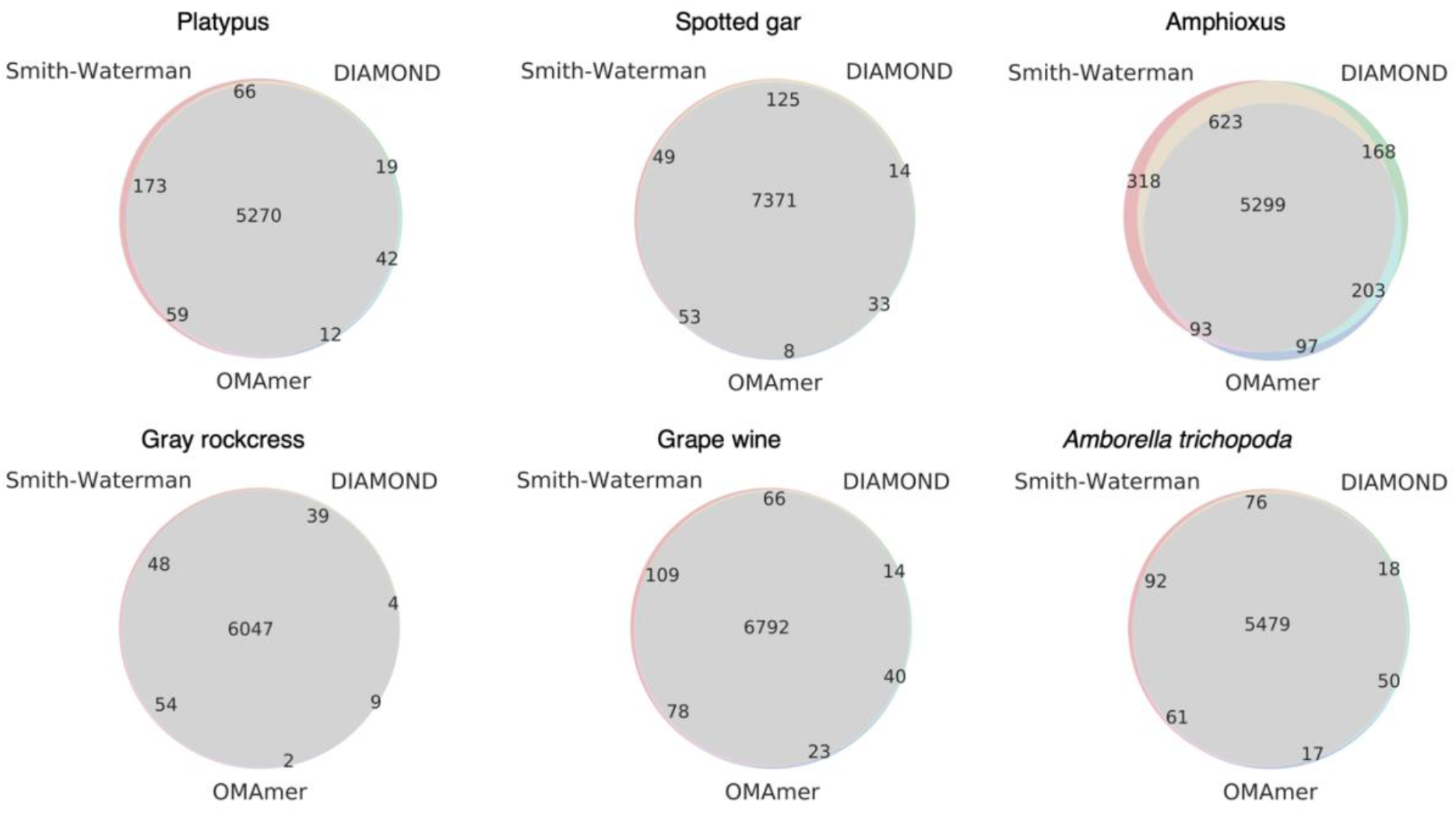
Number of family-level TP queries overlapping between methods. TP sets were defined at F1_max_ for DIAMOND and OMAmer and at the minimum score (1) for Smith-Waterman alignments. These queries were used to assess subfamily assignment.

**Supp. Fig. 2.**
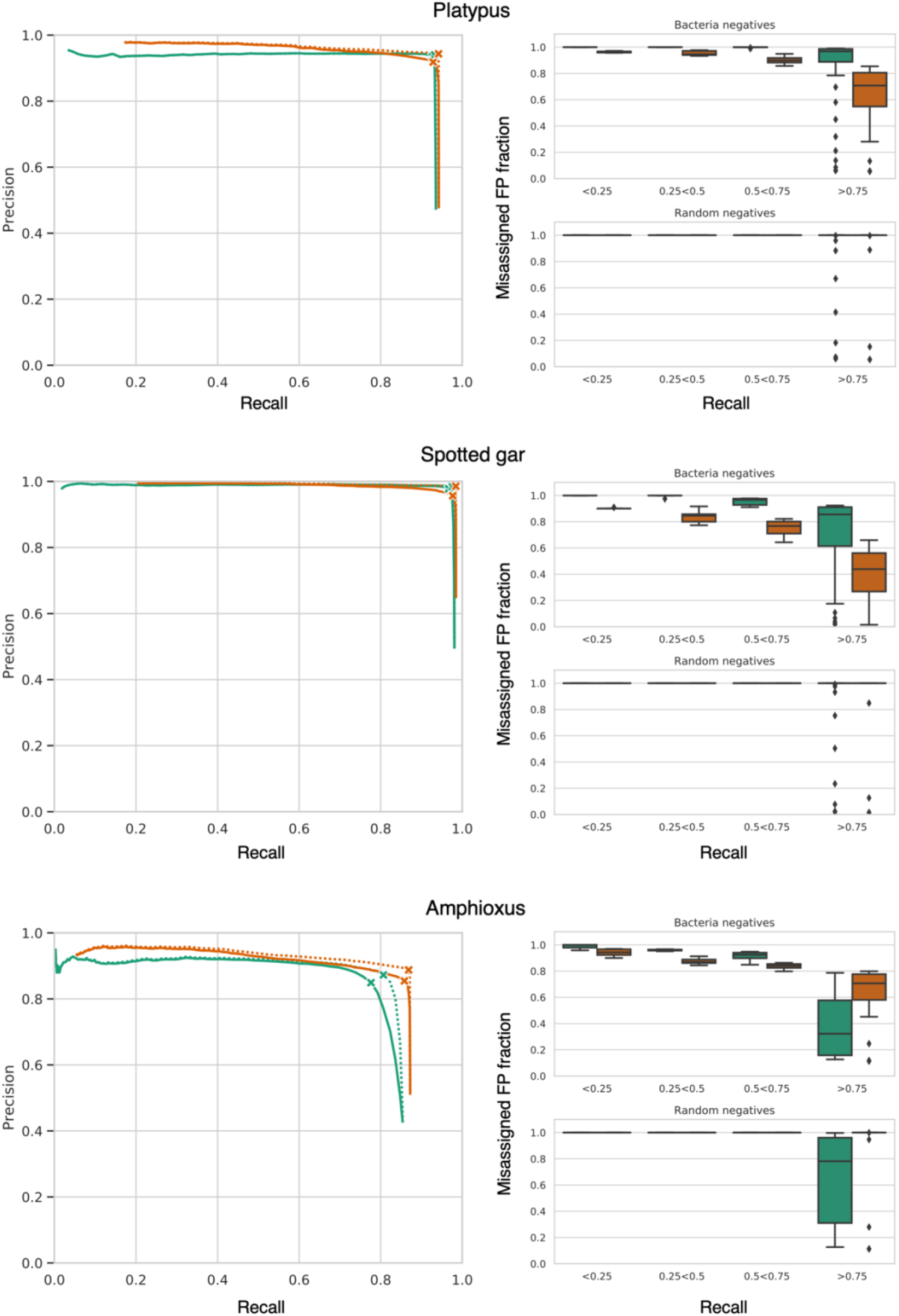
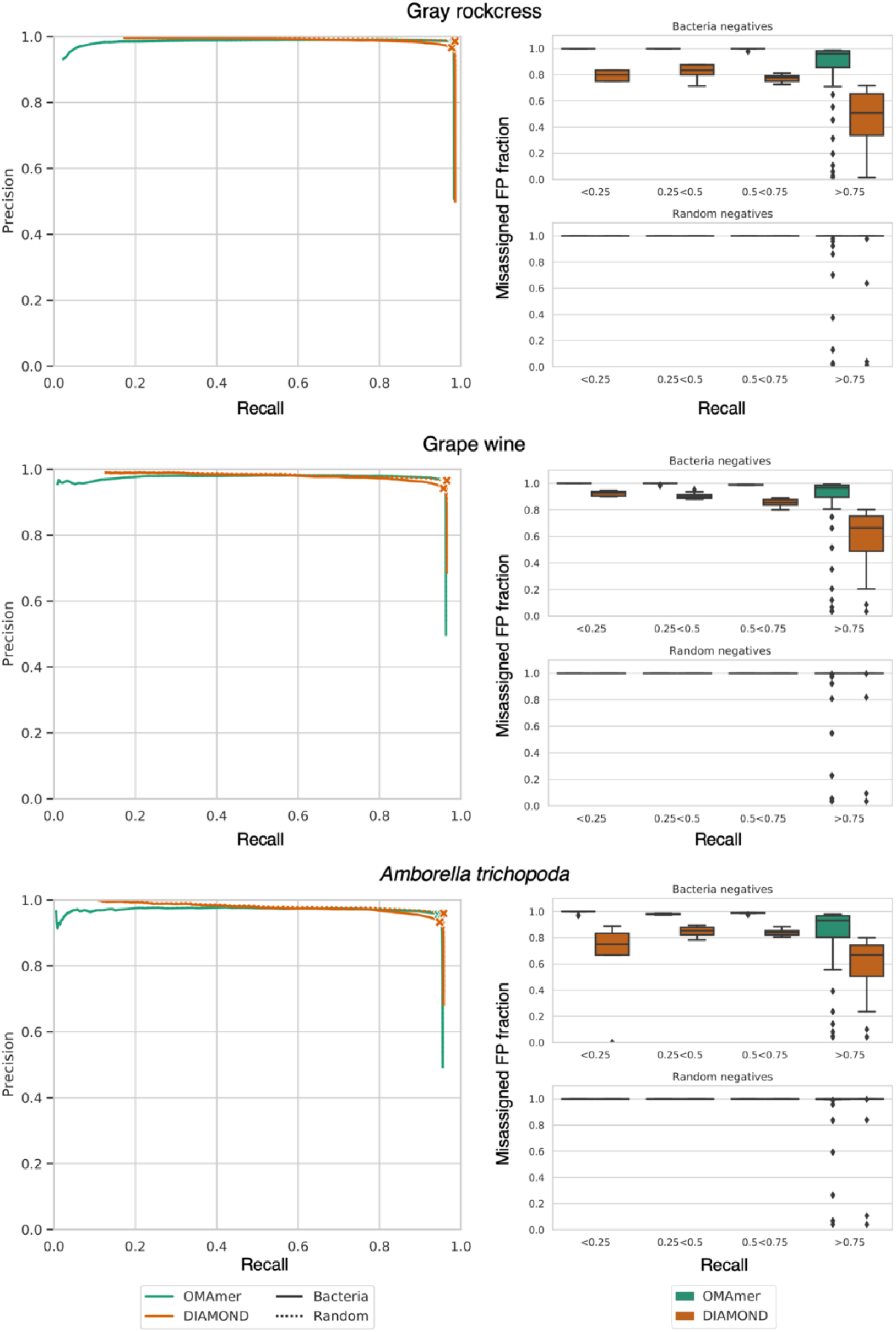
Comparison of family assignments between OMAmer and DIAMOND across negative datasets. (Left) Each curve displays the range of trade-offs between precision and recall when varying the threshold on the OMAmer-score or on the DIAMOND E-value. The curves labeled *Bacteria* refer to analyses using bacteria-specific sequences as negatives whereas those labeled *Random* refer to using random sequences as negatives. Crosses indicate the location of F1_max_ values. (Right) Fraction of FPs coming from the misassignment of positive sequences.

**Supp. Fig. 3.**
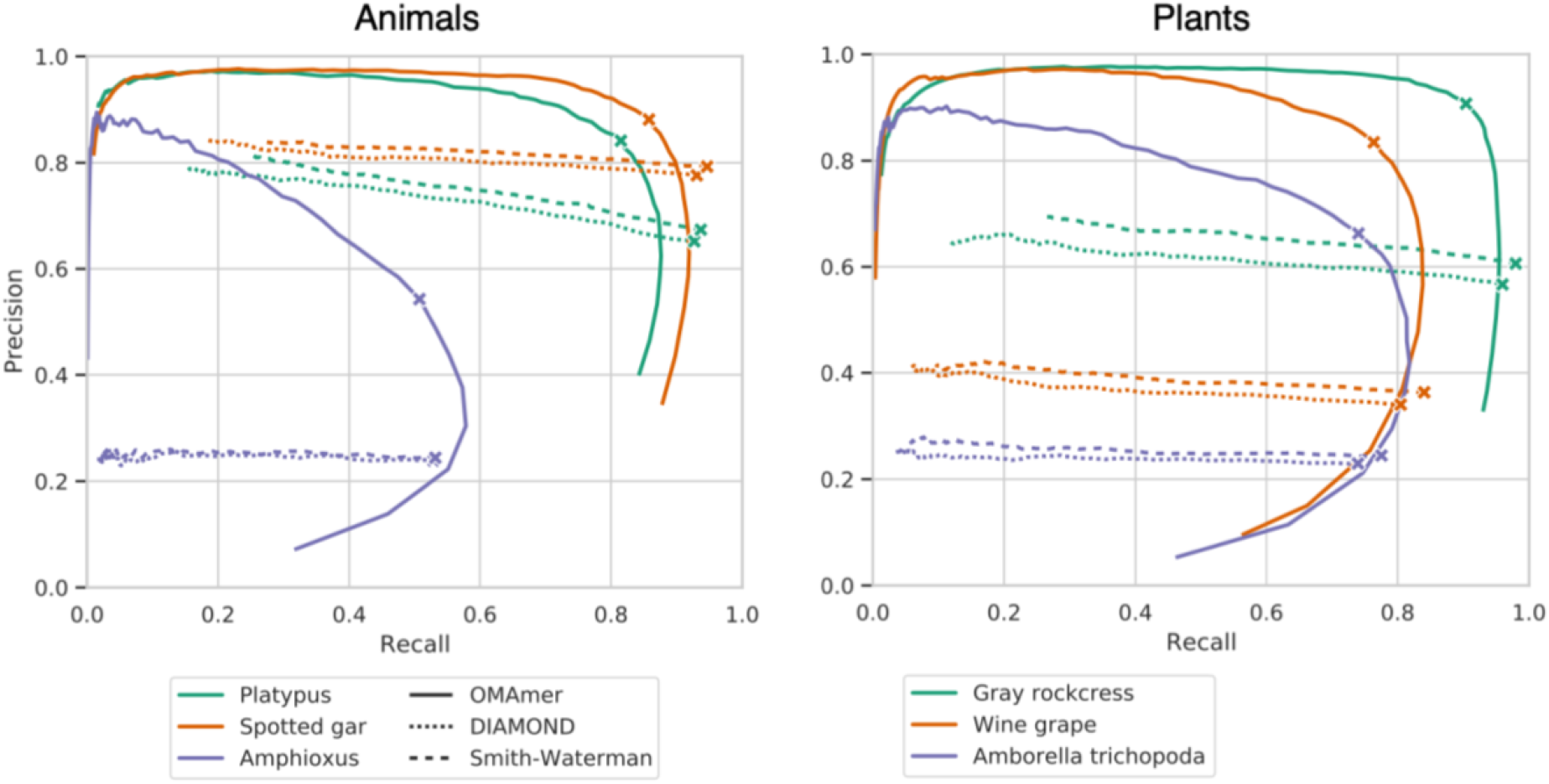
Comparison of subfamily assignments with OMAmer and by closest sequence (Smith-Waterman and DIAMOND). Each curve displays the range of trade-offs between precision and recall when varying the threshold either on the OMAmer-score, on the DIAMOND E-value or on the Smith-Waterman alignment score. These results were computed using the more stringent validation procedure. F1_max_ values are annotated with crosses.

**Supp. Fig. 4.**
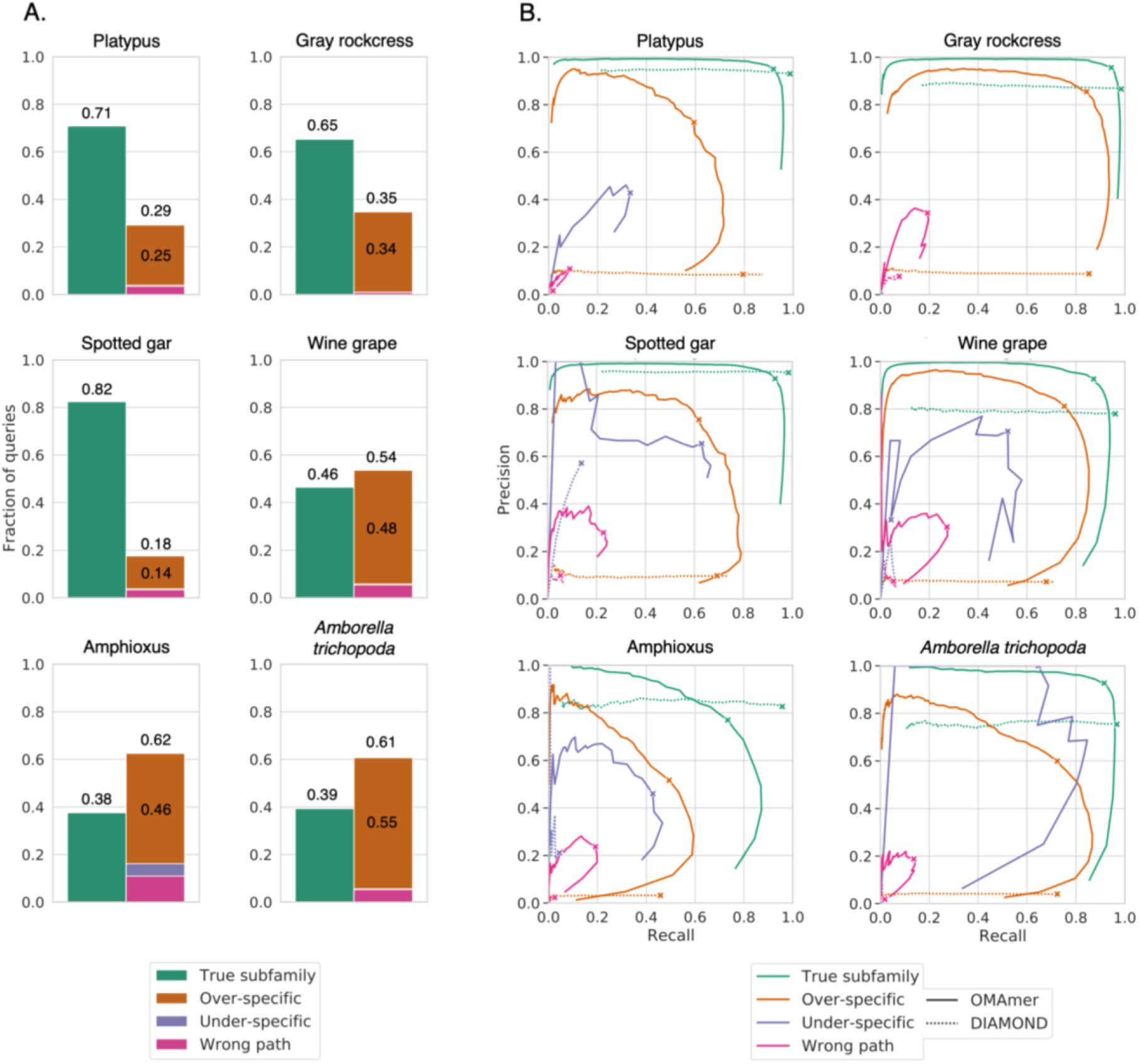
Frequency of closest sequence configurations defined in Fig. 1 and OMAmer accuracy for each. A. The closest sequence to a query was often found in another subfamily. Smith-Waterman alignments were used as proxies for closest sequences. B. These results were computed using the more stringent validation procedure (*See methods).* Each curve displays the range of trade-offs between precision and recall when varying the threshold on the OMAmer-score and on the DIAMOND E-value. They were computed by breaking down queries by closest sequence configurations as in panel A, before the validation procedure itself. F1_max_ values are annotated with crosses. Crosses indicate the location of F1_max_ values. “Over-specific” F1_max_ values are specifically annotated.

**Supp. Fig. 5.**
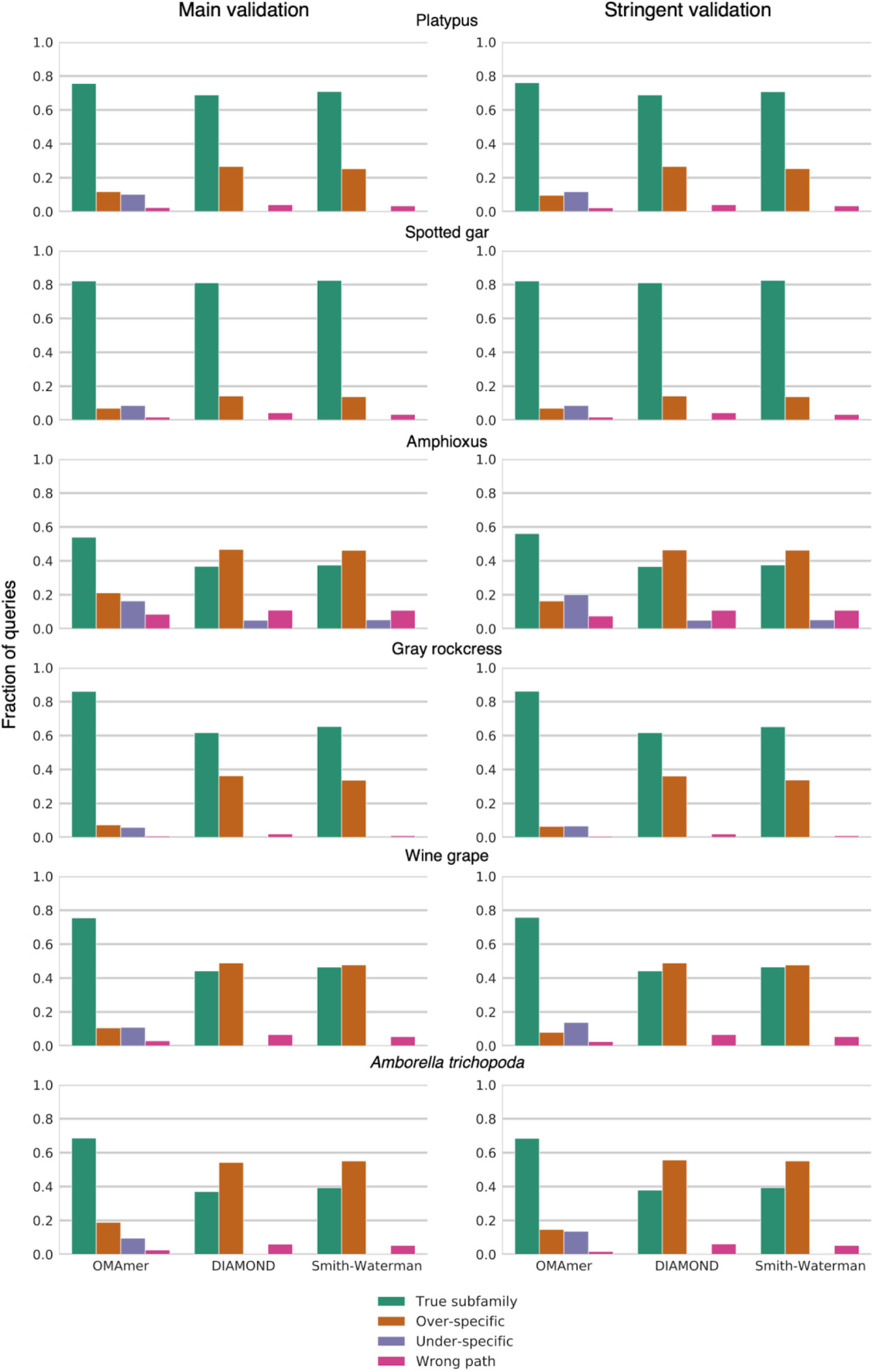
Partitioning of subfamily assignments at F1_max_ into closest sequence configuration.

**Supp. Fig. 6.**
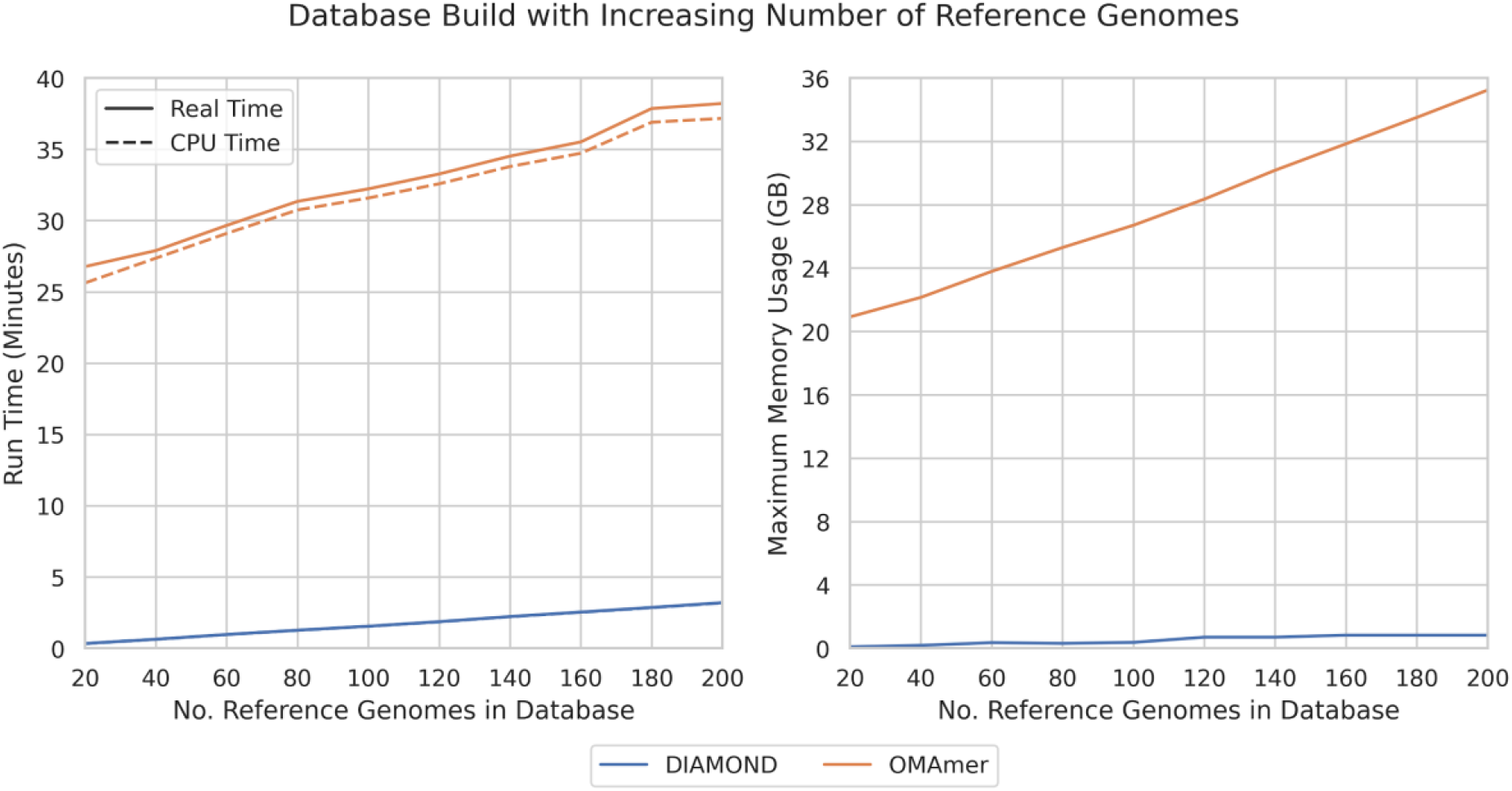
Run time (left) and maximum memory usage (right) during database build for DIAMOND and OMAmer. Whilst OMAmer is slower and requires more memory due to the increased pre-processing to enable fast lookup time, the increase in time and memory is linear with the number of reference genomes in the resulting database.

**Supp. Table 1.**
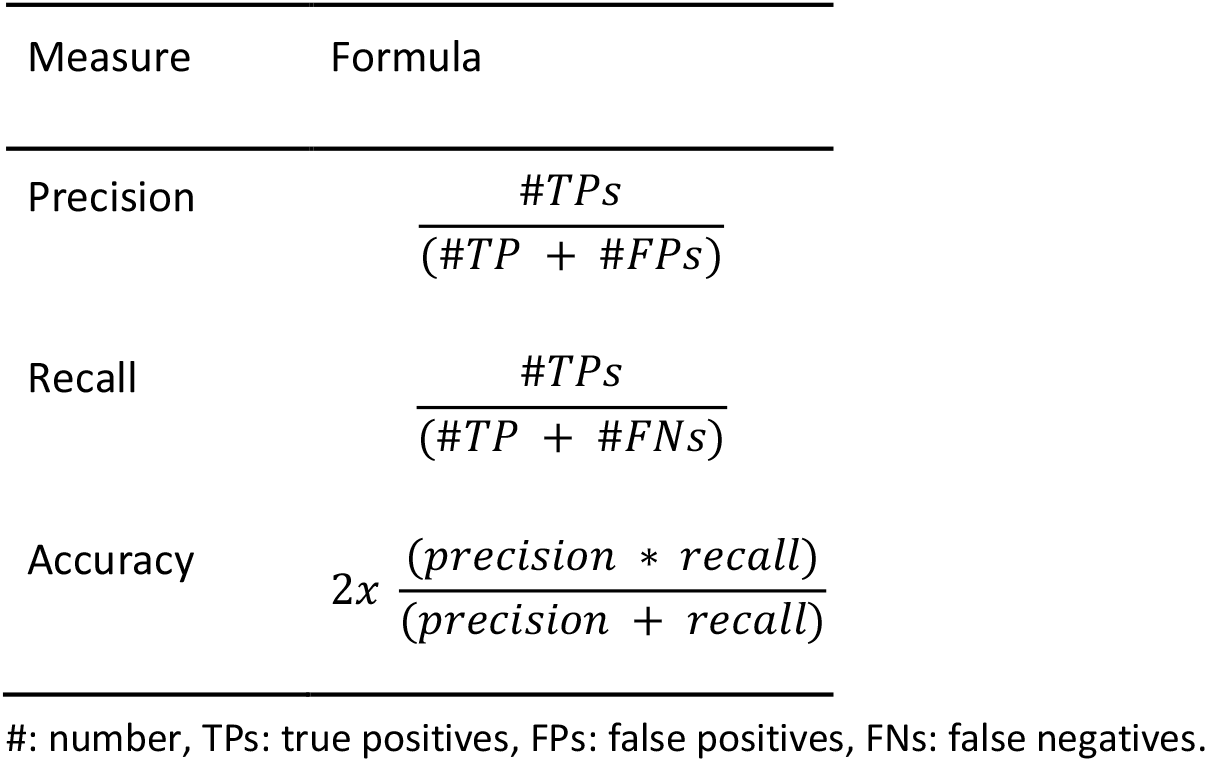
Formulae of validation measures

| Measure | Formula |
| --- | --- |
| Precision | $\frac{\#TPs}{(\#TP + \#FPs)}$ |
| Recall | $\frac{\#TPs}{(\#TP + \#FNs)}$ |
| Accuracy | $2x \frac{(precision * recall)}{(precision + recall)}$ |

#: number, TP: true positives, FP: false positives, FN: false negatives.

**Supp. Table 2.** Species used as queries in benchmarks.

| Species | Scientific name | LCA clade | Divergence age (mya) | Genome scaffold N50 (kb) |
| --- | --- | --- | --- | --- |
| Spotted Gar | *Lepisosteus oculatus* | *Neopterygii* | 320 (Betancur-R *et al.*, 2017) | 6928 (Ensembl LepOcu1 assembly) |
| Platypus | *Ornithorhynchus anatinus* | *Mammalia* | 250 (Upham *et al.*, 2019) | 992 (Ensembl OANA5 assembly) |
| Amphioxus | *Branchiostoma floridae* | *Chordata* | 600 (Peterson and Eernisse, 2016) | 2600 (Putnam *et al.*, 2008) |
| Gray rockcress | *Arabis alpina* | *Brassicaceae* | 27 (Willing *et al.*, 2015) | 788 (Willing *et al.*, 2015) |
| Grape wine | *Vitis vinifera* | *Rosids* | >130 (Jaillon *et al.*, 2007) | 2070 (Jaillon *et al.*, 2007) |
| *Amborella trichopoda* | *Amborella trichopoda* | *Magnoliopsida* | >160 (Amborella Genome Project, 2013) | 4900 (Amborella Genome Project, 2013) |

## References

Altenhoff, A.M. et al. (2013) Inferring hierarchical orthologous groups from orthologous gene pairs. PLoS One, 8, e53786.

Altenhoff, A.M. et al. (2016) Standardized benchmarking in the quest for orthologs. Nat. Methods, 13, 425–430.

Altenhoff, A.M. et al. (2018) The OMA orthology database in 2018: retrieving evolutionaryrelationships among all domains of life through richer web and programmatic interfaces. Nucleic Acids Res., 46, D477–D485.

Altschul, S.F. et al. (1990) Basic local alignment search tool. J. Mol. Biol., 215, 403–410.

Amborella Genome Project (2013) The Amborella genome and the evolution of flowering plants. Science, 342, 1241089.

Barbera, P. et al. (2018) EPA-ng: Massively Parallel Evolutionary Placement of Genetic Sequences. Syst. Biol.

Betancur-R, R. et al. (2017) Phylogenetic classification of bony fishes. BMC Evol. Biol., 17, 162.

Bëinda, K. et al. (2015) Spaced seeds improve k-mer-based metagenomic classification. Bioinformatics, 31, 3584–3592.

Buchfink, B. et al. (2015) Fast and sensitive protein alignment using DIAMOND. Nat. Methods, 12, 59–60.

Conant, G.C. and Wolfe, K.H. (2008) Turning a hobby into a job: how duplicated genes find newfunctions. Nat. Rev. Genet., 9, 938–950.

Dalquen, D.A. et al. (2013) The impact of gene duplication, insertion, deletion, lateral gene transferand sequencing error on orthology inference: a simulation study. PLoS One, 8, e56925.

Ebersberger, I. et al. (2009) HaMStR: profile hidden markov model based search for orthologs in ESTs. BMC Evol. Biol., 9, 157.

Edgar, R.C. (2004) Local homology recognition and distance measures in linear time usingcompressed amino acid alphabets. Nucleic Acids Res., 32, 380–385.

El-Gebali, S. et al. (2019) The Pfam protein families database in 2019. Nucleic Acids Res., 47, D427–D432.

Fox, N.K. et al. (2014) SCOPe: Structural Classification of Proteins--extended, integrating SCOP and ASTRAL data and classification of new structures. Nucleic Acids Res., 42, D304–9.

Gabaldón, T. and Koonin, E.V. (2013) Functional and evolutionary implications of gene orthology. Nat. Rev. Genet., 14, 360–366.

Gladyshev, E.A. et al. (2008) Massive horizontal gene transfer in bdelloid rotifers. Science, 320, 1210–1213.

Glover, N. et al. (2019) Advances and Applications in the Quest for Orthologs. Mol. Biol. Evol., 36, 2157–2164.

Huang, S. et al. (2017) HaploMerger2: rebuilding both haploid sub-assemblies from high-heterozygosity diploid genome assembly. Bioinformatics, 33, 2577–2579.

Huerta-Cepas, J. et al. (2019) eggNOG 5.0: a hierarchical, functionally and phylogenetically annotated orthology resource based on 5090 organisms and 2502 viruses. Nucleic Acids Res., 47, D309–D314.

Huerta-Cepas, J. et al. (2017) Fast Genome-Wide Functional Annotation through Orthology Assignment by eggNOG-Mapper. Mol. Biol. Evol., 34, 2115–2122.

Huson, D.H. et al. (2007) MEGAN analysis of metagenomic data. Genome Res., 17, 377–386.

i5K Consortium (2013) The i5K Initiative: advancing arthropod genomics for knowledge, human health, agriculture, and the environment. J. Hered., 104, 595–600.

Jaillon, O. et al. (2007) The grapevine genome sequence suggests ancestral hexaploidization in major angiosperm phyla. Nature, 449, 463–467.

Kajitani, R. et al. (2019) Platanus-allee is a de novo haplotype assembler enabling a comprehensiveaccess to divergent heterozygous regions. Nat. Commun., 10, 1702.

Koepfli, K.-P. et al. (2015) The Genome 10K Project: a way forward. Annu Rev Anim Biosci, 3, 57–111.

Koski, L.B. and Golding, G.B. (2001) The closest BLAST hit is often not the nearest neighbor. J. Mol. Evol., 52, 540–542.

Kriventseva, E.V. et al. (2019) OrthoDB v10: sampling the diversity of animal, plant, fungal, protist, bacterial and viral genomes for evolutionary and functional annotations of orthologs. Nucleic Acids Res., 47, D807–D811.

Li, L. et al. (2003) OrthoMCL: identification of ortholog groups for eukaryotic genomes. Genome Res., 13, 2178–2189.

Linard, B. et al. (2019) Rapid alignment-free phylogenetic identification of metagenomic sequences. Bioinformatics.

Manber, U. and Myers, G. (1993) Suffix Arrays: A New Method for On-Line String Searches. SIAM J. Comput., 22, 935–948.

Mi, H. et al. (2019) PANTHER version 14: more genomes, a new PANTHER GO-slim and improvements in enrichment analysis tools. Nucleic Acids Res., 47, D419–D426.

Naseeb, S. et al. (2017) Rapid functional and evolutionary changes follow gene duplication in yeast. Proc. Biol. Sci., 284.

Nguyen, N.-P. et al. (2016) HIPPI: highly accurate protein family classification with ensembles of HMMs. BMC Genomics, 17, 765.

Opazo, J.C. et al. (2008) Differential loss of embryonic globin genes during the radiation of placental mammals. Proc. Natl. Acad. Sci. U. S. A., 105, 12950–12955.

Peterson, K.J. and Eernisse, D.J. (2016) The phylogeny, evolutionary developmental biology, and paleobiology of the Deuterostomia: 25 years of new techniques, new discoveries, and new ideas. Org. Divers. Evol., 16, 401–418.

Putnam, N.H. et al. (2008) The amphioxus genome and the evolution of the chordate karyotype. Nature, 453, 1064–1071.

Schreiber, F. et al. (2014) TreeFam v9: a new website, more species and orthology-on-the-fly. Nucleic Acids Res., 42, D922–D925.

Sémon, M. and Wolfe, K.H. (2007) Consequences of genome duplication. Curr. Opin. Genet. Dev., 17, 505–512.

Smith, T.F. and Waterman, M.S. (1981) Identification of common molecular subsequences. J. Mol. Biol., 147, 195–197.

Sonnhammer, E.L.L. and Östlund, G. (2015) InParanoid 8: orthology analysis between 273 proteomes, mostly eukaryotic. Nucleic Acids Res., 43, D234–9.

Steinegger, M. et al. (2019) Protein-level assembly increases protein sequence recovery from metagenomic samples manyfold. Nat. Methods, 16, 603–606.

Tang, H. et al. (2019) TreeGrafter: phylogenetic tree-based annotation of proteins with Gene Ontology terms and other annotations. Bioinformatics, 35, 518–520.

UniProt Consortium (2019) UniProt: a worldwide hub of protein knowledge. Nucleic Acids Res., 47, D506–D515.

Upham, N.S. et al. (2019) Inferring the mammal tree: Species-level sets of phylogenies for questions in ecology, evolution, and conservation. PLoS Biol., 17, e3000494.

Willing, E.-M. et al. (2015) Genome expansion of Arabis alpina linked with retrotransposition and reduced symmetric DNA methylation. Nat Plants, 1, 14023.

Wolf, Y.I. and Koonin, E.V. (2012) A tight link between orthologs and bidirectional best hits in bacterial and archaeal genomes. Genome Biol. Evol., 4, 1286–1294.

Wood, D.E. and Salzberg, S.L. (2014) Kraken: ultrafast metagenomic sequence classification usin gexact alignments. Genome Biol., 15, R46.

Zielezinski, A. et al. (2017) Alignment-free sequence comparison: benefits, applications, and tools. Genome Biol., 18, 186.

Zielezinski, A. et al. (2019) Benchmarking of alignment-free sequence comparison methods. Genome Biol., 20, 144.

